# Clustering and visualization of single-cell RNA-seq data using path metrics

**DOI:** 10.1101/2021.12.14.472627

**Authors:** Andriana Manousidaki, Anna Little, Yuying Xie

## Abstract

Recent advances in single-cell technologies have enabled high-resolution characterization of tissue and cancer compositions. Although numerous tools for dimension reduction and clustering are available for single-cell data analyses, these methods often fail to simultaneously preserve local cluster structure and global data geometry. To address these challenges, we developed a novel analyses framework, <u>S</u>ingle-<u>C</u>ell <u>P</u>ath <u>M</u>etrics <u>P</u>rofiling (scPMP), using power-weighted path metrics, which measure distances between cells in a data-driven way. Unlike Euclidean distance and other commonly used distance metrics, path metrics are density sensitive and respect the underlying data geometry. By combining path metrics with multidimensional scaling, a low dimensional embedding of the data is obtained which preserves both the global data geometry and cluster structure. We evaluate the method both for clustering quality and geometric fidelity, and it outperforms current scRNAseq clustering algorithms on a wide range of benchmarking data sets.

## Introduction

The advance in single-cell RNA-seq (scRNA-seq) technologies in recent years has enabled the simultaneous measurement of gene expression at the single-cell level (1–3). This opens up new possibilities to detect previously unknown cell populations, study cellular development and dynamics, and characterize cell composition within bulk tissues. Despite its similarity with bulk RNAseq data, scRNAseq data tends to have larger variation and larger amounts of missing values due to the low abundance of initial mRNA per cell. To address these challenges, numerous computational algorithms have been proposed focusing on different aspects. Given a collection of single cell transcriptomes from scRNAseq, one of the most common applications is to identify and characterize subpopulations, e.g., cell types or cell states. Numerous clustering approaches have been developed such as *k*-means based methods SC3 (4), SIMLR (5), and RaceID (6); hierarchical clustering based methods CIDR (7), BackSPIN (8), and pcaReduce (9); graph based methods Rphenograph (10), SNN-Cliq (11), Seurat (12), SSNN-Louvain (13), and scanpy (14); and deep-learning based methods scGNN (15), scVI (16), ScDeepCluster (17), DANCE (18), graph-sc (19), GraphSCC (20), scDCC (21), DESC (22),scDHA (23), scziDesk (24),scDSC (25), CELLPLM (26), scDiff (27), scMoGNN (28), scMoFormer (29) and scTAG (30) as summerized in (31).

To visualize and characterize relationships between cell types, it is important to represent it in a low-dimensional space. Many low-dimensional embedding methods have been proposed including UMAP (32), *t*-SNE (33), PHATE (34), and LargeVis (35). However, a key challenge for embedding methods is to simultaneously reduce cluster variance and preserve the global geometry, including the distances between clusters and cluster shapes. For example, Figure 4 illustrates the typical situation on a cell mixture dataset (36): the PCA embedding preserves the global geometry but clusters have high variance; clusters are better separated in the UMAP and *t*-SNE embeddings, but the global geometric structure of the clusters is lost.

**Fig. 1.**
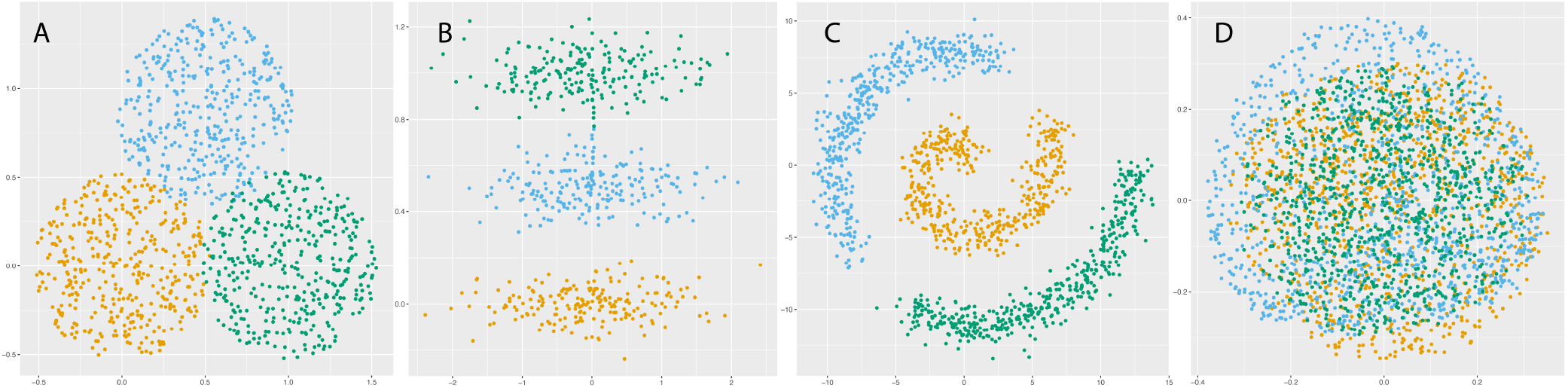
Toy Data Sets: (A) Balls; (B) elongated with bridge; (C) swiss roll; and (D) GL manifold. (A) and (B) show the 2-dimensional data sets. (C) plots the first two coordinates of the Swiss roll. (D) shows the 2-dimensional PCA plot of the SO(3) manifolds.

When choosing a clustering algorithm, there is always an underlying tension between respecting data density and data geometry. Density based methods such as DBSCAN (37, 38) cluster data by connecting together high density regions, regardless of cluster geometry. More traditional approaches such as *k*-means require that clusters are convex and geometrically well separated. However, in many real data, clusters tend to have both nonconvex/elongated geometry and a lack of robust density separation as shown in Figure **??** which consists of three elongated Gaussian distributions and a bridge connecting two of the distributions. The data set is challenging because it exhibits elongated geometry, but methods relying only on density will fail due to the bridge. Such characteristics are commonly observed in scRNA-seq data, especially for cells sampled from a developmental process, as cell types often trace out elongated structures and frequently lack robust density separation. This elongated geometry phenomena is due to the fact that all the cell types originate from stem cells through a trajectory-like differentiation process, and the bridge structures are created by the cells in the transition states. For example, circulating monocytes in the Tabula Muris (TM) lung data set (39) have an elongated cluster structure as illustrated by the PCA plot in Figure 2a, as do the ductal cells in the TM pancreatic data set (see Figure 2c). The UMAP plots of these same data sets illustrate the lack of robust density separation: for TM lung, there is a bridge connecting the alveolar and lung cell types, and also an overlap/bridge between the circulating and invading monocytes (see Figure 2b); for TM pancreatic, the pancreatic A and pancreatic PP cells are not well separated. The combination of elongation and poor density separation make clustering scRNA-seq data sets a challenging task.

**Fig. 2.**
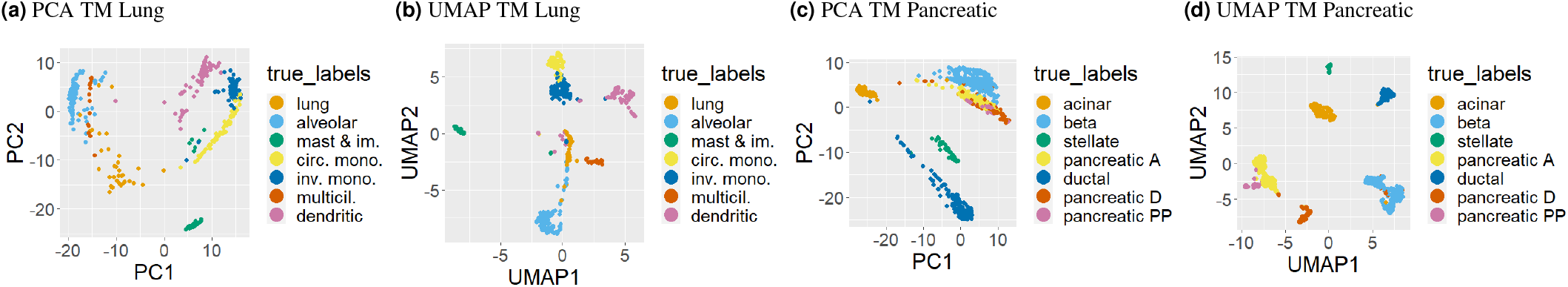
Tabula Muris data sets have elongated clusters in the PCA embedding and clusters connected with a bridge of points in the UMAP embedding. For both PCA and UMAP embeddings, certain clusters are not well-separated and connected by high density regions.

We propose an embedding method based on the *power weighted path metric* which is well suited to this difficult regime. These metrics balance density and geometry considerations in the data via computation of a density-weighted geodesic distance, making them useful for many machine learning tasks such as clustering and semi-supervised learning (40–48). They have performed well in applications such as imaging (46, 47, 49, 50), but their usefulness for the analysis of scRNAseq data remains unexplored.

Because these metrics are density-sensitive, they reduce cluster variance; in addition, these metrics also capture global distance information, and thus preserve global geometry; see Figure 4b. Using the path metric embedding to cluster the data thus yields a clustering method which balances density-based and geometric information.

## Materials and Methods

We first introduce our theoretical framework in Section Path Metrics; The Algorithm Section then describes the details of the proposed scPMP algorithm, and the Assesment Section describes metrics for assessment.

### Path Metrics

We first define a family of power weighted path metrics parametrized by 1 ≤ *p< ∞*.

#### Definition 1

Given a discrete data set *X*, the discrete *p*-power weighted path metric between *a, b* ∈ *X* is defined as

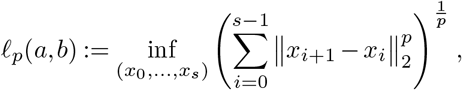

where the infimum is taken over all sequences of points *x*_0_,…, *x*_*s*_ in *X* with *x*_0_ = *a* and *x*_*s*_ = *b*. Note as *p* ⟶ ∞, *l*_*p*_ converges to the γbottleneck edge” distance

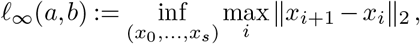

which is well studied in the computer science literature (51– 54). Two points are close in *𝓁*_∞_ if they are connected by a high-density path through the data, regardless of how far apart the points are. On the other hand, when *p* = 1, *𝓁*_1_ reduces to Euclidean distance. If path edges are furthermore restricted to lie in a nearest neighbor graph, *𝓁*_1_ approximates the geodesic distance between the points, i.e. the length of the shortest path lying on the underlying data structure, which is a highly useful metric for manifold learning (55). The parameter *p* governs a trade-off between these two extremes, i.e. it determines how to balance density and geometry considerations when determining which data points should be considered close.

The relationship between *𝓁*_*p*_ and density can be made precise. Assume *n* independent samples from a continuous, nonzero density function *f* supported on a *d*-dimensional, compact Riemannian manifold ℳ (a manifold is a smooth, locally linear surface; see (56)). Then for *p>* 1, *𝓁*_*p*_(*a, b*) converges (after appropriate normalization) to

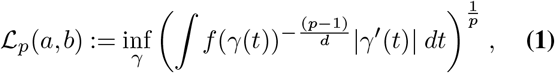

as *n* → ∞, where the infimum is taken over all smooth curves γ : [0, 1] → ℳ connecting *a, b* (57–59). Note |γ^*’*^(*t*)| is simply the arclength element on, ℳ so ℒ_1_ reduces to the standard geodesic distance. When *p* ≠ 1, one obtains a density weighted geodesic distance. The optimal ℒ_*p*_ path is not necessarily the most direct: a detour may be worth it if it allows the path to stay in a high-density region; see Figure 3. Thus the metric is *density-sensitive*, in that distances across high-density regions are smaller than distances across low-density regions; this is a desirable property for many machine learning tasks (60), including trajectory estimation for developmental cells and cancer cells. However the metric is also *geometry preserving*, since it is computed by path integrals on ℳ. The parameter *p* controls the balance of these two properties: when *p* is small, ℒ_*p*_ depends mainly on the geometry of the data, while for large *p*, ℒ_*p*_ is primarily determined by data density.

**Fig. 3.**
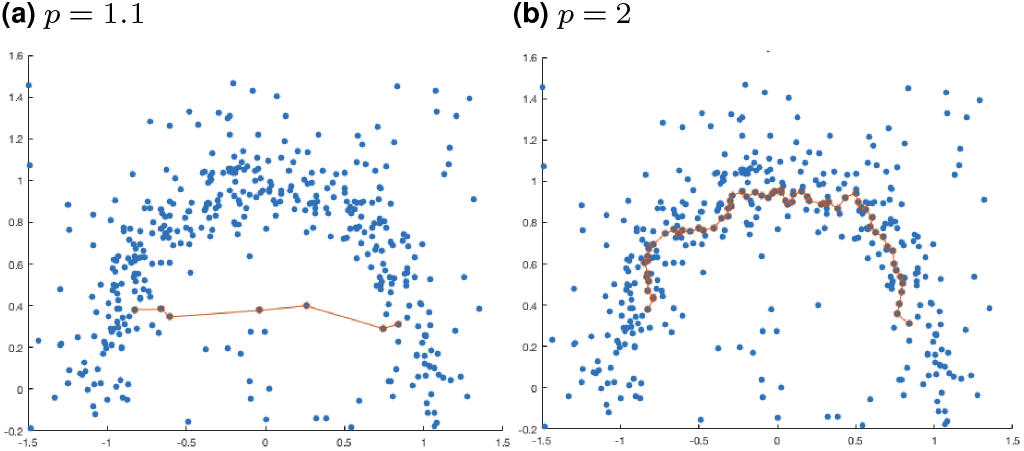
Optimal *𝓁*_*p*_ path between two points in a moon data set.

**Fig. 4.**
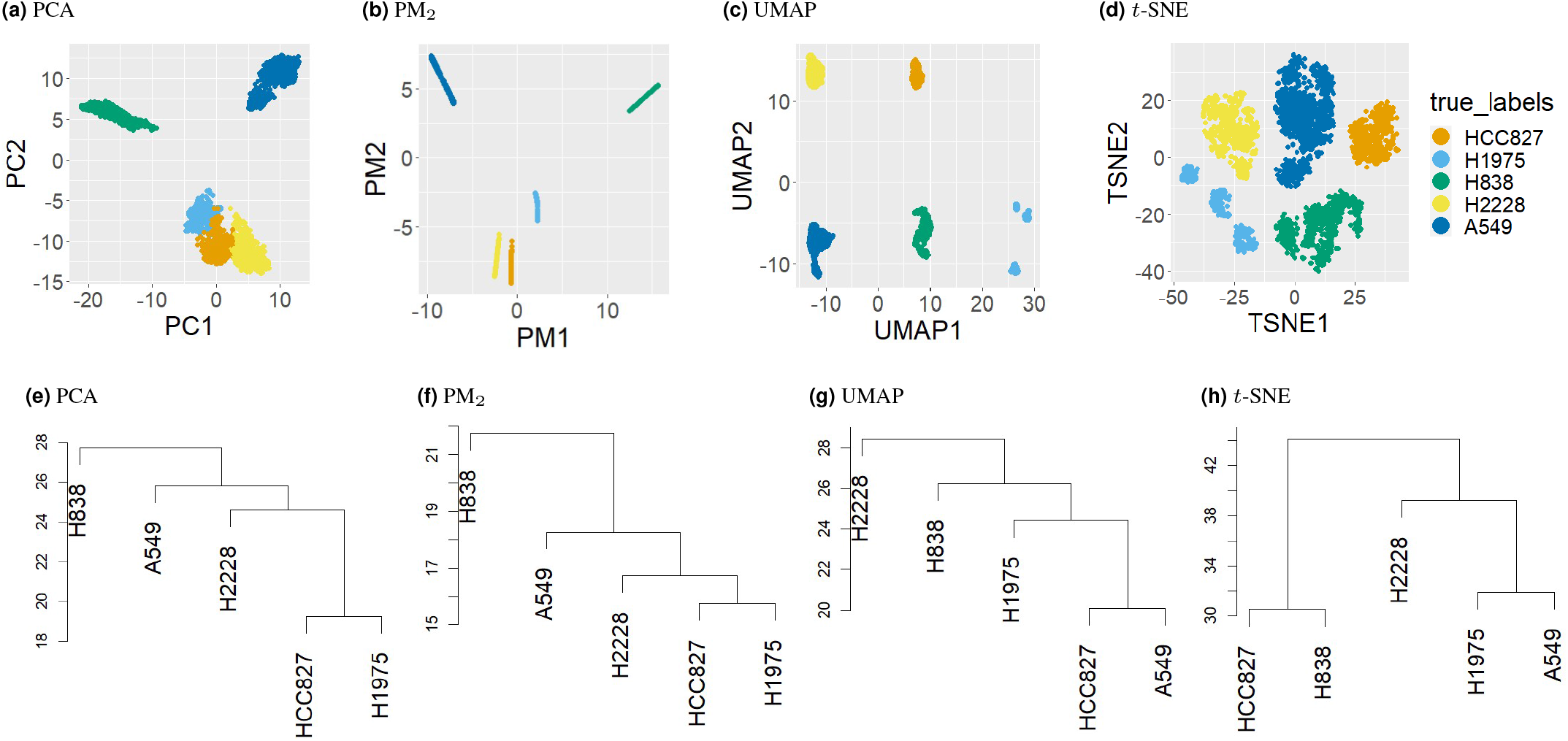
Top row: 2d PCA, PM2, UMAP, and *t*-SNE embeddings of Cell Mix data set, colored by true cell type. Bottom row: average linkage dendrograms of cluster means for the *r*d embeddings, where *r* = 40 for PCA, *r* =4 for PM2 and UMAP, and *r* =3 for *t*-SNE.

Although path metrics are defined in a complete graph, i.e. Definition 1 considers *every* path in the data connecting *a, b*, recent work (46, 61–63) has established that it is sufficient to only consider paths in a *K*-nearest neighbors (*K*NN) graph, as long as *K ≥ C*log*n* for a constant *C* depending on *p, d, f*, and the geometry of the data. By restricting to a *K*NN graph, all pairwise path distances can be computed in *O*(*Kn*^2^) with Dijkstra’s algorithm (64).

**Algorithm**. We consider a noisy data set of *n* data points 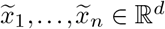, which form the rows of noisy data matrix 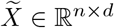. We first denoise the data with a local averaging procedure, which has been shown to be advantageous for manifold plus noise data models (65) and contributes to the improvement of clustering performance on scRNAseq data sets as explored in Supplementary Note 1. More specifically, we replace 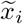 with its local average:

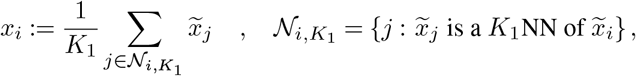

and let *X ∈* ℝ^*n×d*^ denote the denoised data matrix.

#### Algorithm 1

scPMP

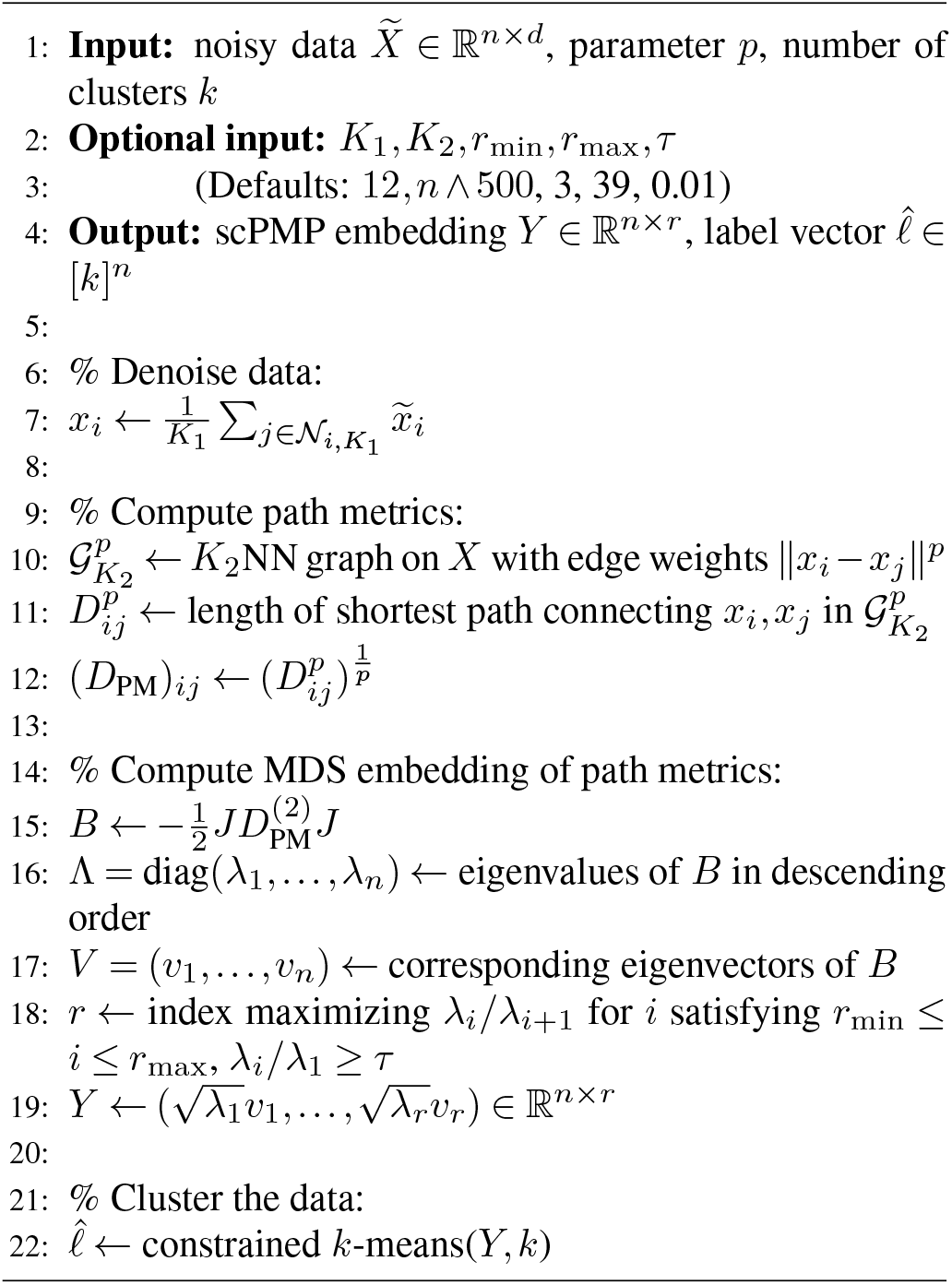

We then fix *p* and compute the *p*-power weighted path distance between all points in *X* to obtain pairwise distance matrix *D*_PM_ ∈ ℝ^*n×n*^. More precisely, we let 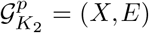 be the graph on *X* where *x*_*i*_, *x*_*j*_ are connected with edge weight 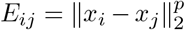 if *x*_*i*_ is a *K*_2_NN of *x*_*j*_ or *x*_*j*_ is a *K*_2_NN of *x*_*i*_. We then compute 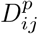 as the total length of the shortest path connecting *x*_*i*_, *x*_*j*_ in 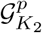, and define *D*_PM_ by 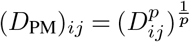.

We next apply classical multidimensional scaling (66) to obtain a low-dimensional embedding which preserves the path metrics. Specifically, we define the path metric MDS matrix 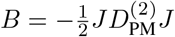 where 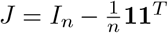 is the centering matrix, **1** ∈ ℝ^*n*^ is a vector of all 1’s, and 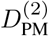 is obtained from *D*_PM_ by squaring all entries. We let the spectral decomposition of *B* be denoted by *B* = V ΛV ^*T*^, where Λ = diag(_1_,…, *λ*_*n*_), *V* = (*v*_1_,…, *v*_*n*_) ∈ ℝ^*n×n*^ contain the eigenvalues and eigenvectors of *B* in descending order. The embedding dimension *r* is then chosen as the index *i* which maximizes the eigenratio *λ*_*i*_*/λ*_*i*+1_ (67), with the following restrictions: we constrain 3 ≤*i ≤*39 and only consider ratios *λ*_*i*+1_*/λ*_*i*_ between γlarge” eigenvalues, i.e. we require *λ*_*i*_*/λ*_1_≥ 0.01. The scPMP embedding is then defined by 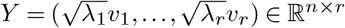.

Finally, we apply *k*-means to the scPMP embedding to obtain cluster labels. Specifically, we let 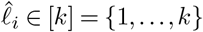 be the cluster label of *x*_*i*_ returned by running *k*-means on *Y* with *k* clusters and 20 replicates. Since *k*-means may return highly imbalanced clusters, cluster sample sizes were constrained to be at least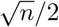. Specifically, if *k*-means returned a tiny cluster, *k* was increased to *k* + 1, and the tiny cluster merged with the closest non-trivial cluster. This entire procedure is summarized in the pseudocode in Algorithm 1.

We note that the computational bottleneck for scPMP is the computation and storage of all pairwise path distances, which has complexity *O*(*n*^2^ log *n*) when *K*_2_ = *O*(log *n*). However this quadratic cost can be avoided by utilizing a low rank approximation of the squared distance matrix via the Nystrom method (68–72). For example, (73) propose a fast, quasi-linear implementation of MDS which only requires the computation of path distances from a set of *q* landmarks, so that the complexity of computing path distances is reduced to *O*(*qn* log *n*). Our implementation of scPMP includes the option to use this landmark-based approximation and is thus highly scalable.

We also note that an important consideration in the fully unsupervised setting is how to select the number of clusters *k*. This is a rather ill-posed question with multiple reasonable answers due to hierarchical cluster structure. We do not focus on this in the current article, and scPMP assumes the number of clusters is given. However we emphasize that when *k* is unknown, the scPMP embedding offers a useful tool for selecting a reasonable number of clusters. For example, Line 21 of Algorithm 1 can be repeated for a range of candidate *k* values to obtain candidate clusterings 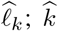 can then be chosen so that 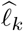 optimizes a cluster validity criterion such as the silhouette criterion (74, 75). Alternatively, one could build a graph with distances computed in the scPMP embedding, and estimate *k* as the number of small eigenvalues of a corresponding graph Laplacian (47, 76).

### Assessment

We evaluate the performance of scPMP with respect to (1) cluster quality and (2) geometric fidelity on a collection of labeled benchmarking data sets with ground truth labels *𝓁*. There are many helpful metrics for the quality of the estimated cluster labels 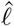, and we compute the adjusted rand index (ARI), entropy of cluster accuracy (ECA), and entropy of cluster purity (ECP). Definitions of ECA and ECP can be found in Supplementary Note 2. We compare our clustering results with the output of *k*-means, DBSCAN (37, 38), *k*-means on *t*-SNE embedding (33), DBSCAN on UMAP embedding (32) and for scRNAseq data sets additionally with the following scRNAseq clustering methods: SC3 (4), scanpy (14), RaceID3 (77), SIMRL (5) and Seurat (12). Assessing the geometric fidelity of the low-dimensional embedding *Y* is more delicate; we want to assess whether the embedding procedure preserves the global relative distances between clusters. We first compute the mean of each cluster as in (33) using the ground truth labels, i.e.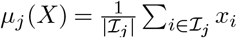 where ℐ_*j*_ = {*i* : *𝓁*_*i*_ = *j*}; we then define 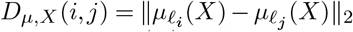. Similarly, we compute the means *μ*_*j*_(*Y*) in the scPMP embedding, and define 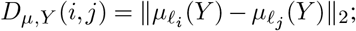 we then compare *D*_*μ,X*_ and *D*_*μ,Y*_. Specifically, we define the geometric perturbation *π* by:

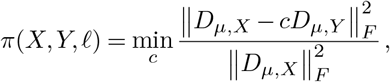

where ∥ ·∥_*F*_ is the Frobenius norm. The *c* achieving the minimum is easy to compute, and one obtains

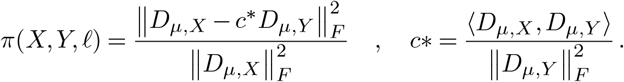

We compare *π*(*X, Y, 𝓁*) with the geometric perturbation of other embedding schemes for *X*, i.e. with *π*(*X, U, 𝓁*) for *U* equal to the UMAP (32) and *t*-SNE (33) embeddings. Note that *π* is not always a useful measure: for example if *X* consisted of concentric spheres sharing the same center, the metric would be meaningless, as the distance between cluster means would be zero. Nevertheless, in most cases *π* is a helpful metric for quantifying the preservation of global cluster geometry.

## Results

We apply scPMP to both a collection of toy manifold data sets and a collection of scRNAseq data sets. Results are reported in Sections and respectively. The default parameter values reported in scPMP were used on all data sets.

### Manifold Data

We apply scPMP for *p* = 1.5, 2, 4 to the following four manifold data sets:

- **Balls** (*n* = 1200, *d* = 2, *k* = 3): Clusters were created by uniform sampling of 3 overlapping balls in ℝ^2^; see Figure 1A.
- **Elongated with bridge** (denoted EWB, *n* = 620, *d* = 2, *k* = 3): Clusters were created by sampling from 3 elongated Gaussian distributions. A bridge was added connecting two of the Gaussians; see Figure 1B.
- **Swiss roll** (*n* = 1275, *d* = 3, *k* = 3): Clusters were created by uniform sampling from three distinct regions of a Swiss roll; 3-dimensional isotropic Gaussian noise (*σ* = 0.75) was then added to the data. Figure 1C shows the first two data coordinates.
- **SO(3) manifolds** (*n* = 3000, *d* = 1000, *k* = 3): For 1 ≤ *i ≤* 3, the 3-dimensional manifold ℳ_*i*_ ⊆ ℝ^9^ is defined by fixing three eigenvalues *D*_*i*_ = diag(*λ*_1_, *λ*_2_, *λ*_3_) and then defining _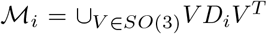_, where SO(3) is the special orthogonal group. After fixing *D*_*i*_, we randomly sample from ℳ _*i*_ by taking random orthonormal bases *V* of ℝ^3^. A noisy, high-dimensional embedding was then obtained by adding uniform random noise with standard deviation *σ* = 0.0075 in 1000 dimensions. Figure 1D shows the first two principal components of the data, which exhibits no cluster separation.

The data sets were chosen to illustrate various cluster separability characteristics. For the balls, the clusters have good geometric separation but are not separable by density. For the Swiss roll and SO(3), the clusters have a complex and inter-twined geometry but are well separated in terms of density. For EWB, clusters are both elongated and lack robust density separability due to the bridge, and one expects that methods which rely too heavily on either geometry or density will fail. The ARIs achieved by scPMP, *k*-means based methods, DBSCAN based methods, and Seurat are reported in Table 2. See Tables 3 and 4 in Supplementary Note 2 for ECP and ECA. As expected, *k*-means out performs all methods on the balls but performs very poorly on all other data sets. DBSCAN and Seurat achieve perfect accuracy on the Swiss roll and SO(3) but perform rather poorly on the balls and EWB, although Seurat does noticeably better than DBSCAN. scPMP with *p* = 2 (PM_2_), is the only method which achieves a high ARI (*>* 90%) and a low ECP and ECA (*<* 0.15) on all data sets.

**Table 1.** Notations.

| Notation | Definition |
| --- | --- |
| ARI | Adjusted Rand Index |
| ECP | Entropy of Cluster Purity |
| ECA | Entropy of Cluster Accuracy |
| UMAP | Uniform Manifold Approximation and Projection |
| t-SNE | t-distributed Stochastic Neighbor Embedding |
| PCA | Principal Component Analysis |
| MDS | Mutlidimensional Scaling |
| scPMP | Single-Cell Path Metrics Profiling |
| PM | Path Metrics |
| $p$ | Parameter of power weighted path metrics |
| $PM_p$ | scPMP clustering with path metric parameter $p$ |
| NN | Nearest Neighbors |
| $K_1$ | Number of NN in local averaging |
| $K_2$ | Number of NN for path metric distance |
| $k$ | Number of clusters |
| $d$ | Number of features of data set |
| $n$ | Number of samples |
| $f$ | Density function of samples |
| $\pi$ | Geometric pertrubations |

**Table 2.**
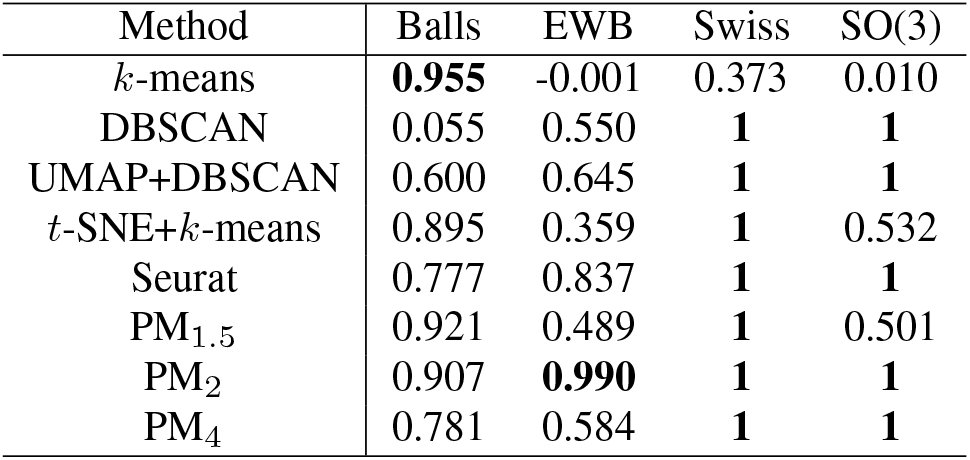
The results of clustering accuracy (ARI) for manifold data.

**Table 3.** Geometric perturbation for manifold data. The *r*d UMAP embeddings were computed with an embedding dimension of *r* =5 for the balls, EWB, Swiss roll and *r* = 7 for SO(3), which corresponded to the estimated dimension for both PM_1.5_ and PM_2_. For *t*-SNE, *r* =3 for all data sets.

| Method | Balls | EWB | Swiss | SO(3) |
| --- | --- | --- | --- | --- |
| 2d UMAP | 0.001 | 0.006 | 0.305 | 0.071 |
| $rd$ UMAP | **0** | 0.033 | 0.339 | **0.054** |
| 2d $t$-SNE | **0** | **0.004** | 0.187 | 0.171 |
| $rd$ $t$-SNE | **0** | 0.042 | 0.074 | 0.157 |
| 2d PM<sub>1.5</sub> | **0** | 0.033 | **0.002** | 0.103 |
| $rd$ PM<sub>1.5</sub> | **0** | 0.023 | 0.011 | 0.154 |
| 2d PM<sub>2</sub> | **0** | 0.146 | 0.025 | 0.156 |
| $rd$ PM<sub>2</sub> | **0** | 0.068 | 0.025 | 0.179 |
| 2d PM<sub>4</sub> | 0.003 | 0.191 | 0.056 | 0.194 |
| $rd$ PM<sub>4</sub> | 0.004 | 0.157 | 0.056 | 0.194 |

**Table 4.** The results of clustering accuracy (ARI) for scRNAseq data.

| Method | RNA1 | RNA2 | TMLung | Beta | TMPanc | BaronPanc | PBMC4K | CellMix |
| --- | --- | --- | --- | --- | --- | --- | --- | --- |
| SC3 | 0.637 | 0.827 | 0.798 | 0.969 | 0.894 | 0.767 | 0.889 | 1 |
| scanpy | 0.620 | 0.825 | 0.796 | 0.910 | 0.615 | 0.966 | 0.977 | 1 |
| RaceID3 | 0.730 | 0.520 | **0.900** | 0.714 | 0.751 | 0.651 | 0.763 | 1 |
| SIMLR | 0.878 | 0.792 | 0.727 | **0.975** | 0.599 | 0.698 | 0.705 | 1 |
| Seurat | 0.792 | 0.667 | 0.843 | 0.891 | 0.547 | 0.941 | 0.889 | 0.993 |
| Seurat_def | 0.714 | 0.785 | 0.764 | 0.919 | 0.798 | **0.971** | 0.975 | 1 |
| $k$-means | 0.921 | 0.786 | 0.848 | 0.969 | 0.840 | 0.662 | 0.747 | 1 |
| DBSCAN | **0.952** | 0.826 | 0.587 | 0.568 | 0.734 | 0.724 | 0.889 | 1 |
| UMAP+DBSCAN | 0.926 | 0.892 | 0.619 | 0.565 | 0.893 | 0.848 | 0.974 | 1 |
| $t$-SNE+$k$-means | 0.943 | 0.915 | 0.753 | 0.969 | 0.620 | 0.641 | 0.596 | 0.878 |
| $PM_{1.5}$ | 0.939 | 0.924 | 0.888 | 0.969 | 0.626 | 0.804 | 0.754 | 1 |
| $PM_2$ | 0.939 | **0.973** | 0.808 | 0.969 | **0.918** | 0.969 | 0.757 | 1 |
| $PM_4$ | 0.939 | 0.939 | 0.731 | 0.921 | 0.775 | 0.853 | **0.978** | 1 |

Table 3 reports the geometric perturbation of the embedding produced by scPMP and compares with UMAP and *t*-SNE. Since scPMP generally selects an embedding dimension *r>* 2, to ensure a fair comparison the geometric perturbation was computed in both the 2d and *r*-dimensional (*r*d) embeddings for all methods, where for UMAP *r* is the dimension selected by Algorithm 1 and for *t*-SNE *r* =3 (note *r ≤*3 was required in Rtsne implementation). Overall PM_1.5_ achieved the lowest geometric perturbation, although all methods had small perturbation on the Balls data set and *t*-SNE had the lowest perturbation on EWB. We point out however that for both the Swiss roll and SO(3), the metric may not be meaningful due to the complicated cluster geometry.

### scRNAseq Data

We apply scPMP for *p* = 1.5, 2, 4 to the following synthetic scRNAseq data sets:

- **RNA mixture:** Benchmarking scRNAseq data set from (36). RNAmix1 was processed with CEL-seq2 and has *n* = 296 cells and *d* = 14687 genes. RNAmix2 was processed with Sort-seq and has *n* = 340 cells and *d* = 14224 genes. For the creation of the two data sets, RNA was extracted in bulk for each of the following cell lines: H2228, H1975, HCC827. Then the RNA was mixed in *k* = 7 different proportions (each defining a ground truth cluster label), diluted to single cell equivalent amounts ranging from 3.75pg to 30pg, and processed using CEL-seq2 and SORT-seq. See here for Supplemental info including ground truth geometric structure.
- **Simulated beta:** Simulated data set of *n* = 473 beta cells and *d* = 2279 genes, created based on SAVER (78) and scImpute (79). First, we subset the Baron’s Pancreatic data set (80) to include only Beta cells. As in (79), we randomly choose 10% of the genes to operate as marker genes. Then, we split the cells to *k* =3 clusters and each cluster is assigned a different group of marker genes. For each cluster we scale up the mean expression of its marker genes. Lastly, to simulate the drop out effect, as in (78), we multiply each cell by an efficiency loss constant drawn by Gamma(10, 100). Using *S* to refer to the data matrix resulting from the above steps, the final simulated data *X* is obtained by letting *X*_*ij*_ be drawn from Poisson(*S*_*ij*_).

In addition to the synthetic data, we evaluate the performance of scPMP on the following real scRNAseq data sets:

- **Cell mixture data set:** Another benchmarking data set from (36) consisting of a mixture of *k* = 5 cell lines created with 10x sequencing platform. The cell line identity of a cell is also its true cluster label. The data set consists of *n* = 3822 cells and *d* = 11786 genes; we removed multiplets, based on the provided meta-data file and kept 3000 most variable genes after SCT tranformation (81, 82).
- **Baron’s pancreatic:** Human pancreatic data set generated by (80). After quality control and SAVER imputation, there are *d* = 14738 genes and *n* = 1844 cells. For analysis purposes cells that belong in a group with less than 70 members were filtered out to reduce to *k* = 8 cell types. Also, we kept only the 3000 most variable genes after SCT tranformation (81, 82). The cell types associated with each cell were obtained by an iterative hierarchical clustering method that restricts genes enriched in one cell type from being used to separate other cell types. The enriched markers in every cluster defined the cell type of the cells that belong in that cluster.
- **Tabula Muris data sets:** Mouse scRNAseq data for different tissues and organs (39). We select the pancreatic data (TM Panc) with *n* = 1444 cells and *d* = 23433 genes and the lung data (TM Lung) with *n* = 453 cells and *d* = 23433 genes. Both data sets have *k* =7 different cell types which were characterized by an FACS-based full length transcript analysis.
- **PBMC4k data set:** This data set includes the gene expression of Peripheral Blood Mononuclear Cells. The raw data are available from 10X Genomics at https://support.10xgenomics.com/single-cell-gene-expression/datasets/2.1.0/pbmc4k. After quality control, saver imputation, and removing the two smallest cell types, there are *d* = 16655 genes and *n* = 4316 cells in the dataset. Also, we merge CD8+ T-cells and CD4+ T-cells in one type named T-cells resulting in *k* = 4 cell types. The ground truth cell types are provided by SingleR annotation after marker gene verification in github.com/SingleR.

Details about the pre-processing of data sets can be found in Supplementary Note 1. For the following UMAP and *t*-SNE results, Linnorm normalization was applied without denoising, as this normalization gave the best results. Note Seurat_def refers to the results of the entire Seurat pipeline, whereas Seurat refers to the result of using Seurat clustering on data with the same processing and normalization as for PM. The embedding dimension *r* selected by scPMP ranged from 3 to 7 for PM_1.5_ and PM_2_, and from 3 to 11 for PM_4_.

Table 4 reports the clustering accuracy regarding ARI achieved by scPMP and other methods; see Tables 5 and 7 in Supplementary Note 2 for ECP and ECA. The path metric methods perform equally well or better than the rest of the methods. Once again PM_2_ exhibits the best overall performance, with a high ARI (≥90%) on all data sets except TM lung and PBMC4K; the next best method is PM_4_, which achieves a high ARI on all but 3 data sets. Seurat_def and PM_1.5_ had a low ARI for 4 of 8 data sets; scanpy, *k*-means, UMAP+DBSCAN and *t*-SNE+*k*-means had a low ARI on 5 of the 8 data sets; SC3, RaceId3, SIMLR and Seurat had a low ARI (*<* 90%) on 6 of the 8 data sets. These results indicate that incorporating both density-based and geometric information when determining similarity generally leads to more robust results for scRNA-seq data. Moreover, PM_2_ achieves the best median ECP and median ECA values across all RNA data sets. Although the optimal balance depends on the data set (for example PBMC4K does best with *p* = 4, while TMLung does best with *p* = 1.5), path metrics with a moderate *p* exhibit the best performance across a wide range of data sets.

**Table 5.** Geometric perturbation for RNA data. For *r*d UMAP *r* = 7, 6, 5, 3, 5, 9, 3, 4 for cximum of the PM_1.5_ dimension and the PM_2_ dimension. For *r*d *t*-SNE *r* = 3.

| Method | RNA1 | RNA2 | TMLung | Beta | TMPanc | BaronPanc | PBMC4k | CellMix |
| --- | --- | --- | --- | --- | --- | --- | --- | --- |
| 2d UMAP | 0.122 | 0.142 | 0.057 | 0.025 | 0.064 | 0.115 | 0.015 | 0.090 |
| rd UMAP | 0.160 | 0.131 | 0.092 | 0.026 | 0.036 | 0.129 | 0.027 | 0.050 |
| 2d $t$-SNE | 0.059 | 0.054 | 0.042 | 0.024 | 0.048 | 0.206 | 0.038 | 0.061 |
| rd $t$-SNE | 0.035 | 0.054 | 0.027 | 0.016 | 0.040 | 0.229 | 0.050 | 0.033 |
| 2d $PM_{1.5}$ | **0.010** | 0.013 | 0.046 | 0.002 | 0.076 | 0.067 | 0.028 | 0.098 |
| rd $PM_{1.5}$ | 0.017 | **0.009** | **0.006** | **0** | **0.019** | **0.006** | **0.007** | **0.007** |
| 2d $PM_2$ | 0.040 | 0.040 | 0.085 | 0.003 | 0.150 | 0.103 | 0.050 | 0.101 |
| rd $PM_2$ | 0.048 | 0.036 | 0.029 | 0.003 | 0.051 | 0.010 | 0.013 | 0.008 |
| 2d $PM_4$ | 0.108 | 0.135 | 0.246 | 0.016 | 0.265 | 0.193 | 0.069 | 0.107 |
| rd $PM_4$ | 0.100 | 0.082 | 0.083 | 0.015 | 0.099 | 0.027 | 0.029 | 0.008 |

For BaronPanc, we observe that Seurat_def achieves a slightly higher ARI than all the reported path metric methods (*p* = 1.5, 2, 4). However, a significant advantage of scPMP over Seurat is the high clustering performance on a wide range of sample sizes. To demonstrate our claim we compare the ARI results in different down-sampled versions of BaronsPanc. We selected a stratified sample of 50%, 25% and 10% of the cells of the BaronPanc data set. The results can be found in Table 8 of Supplementary Note 2. We observed no ARI deterioration for scPMP for the 50% and 25% down-sampled data set and only a moderate decrease for the 10% down-sampled dataset (ARI of 0.67 at 10% downsampling for *p* = 1.5). On the contrary, there is significant ARI deterioration both for Seurat and Seurat_def; in particular, at 10% downsampling the ARI deteriorates to 0.405 for Seurat and to 0.185 for Seurat_def. Notice that in the 10% downsampled data set, we use regular *k*-means for PM_2_ to allow for the prediction of smaller sized clusters.

We also investigated whether we could learn the ground truth number of clusters by optimizing the silhouette criterion in the scPMP embedding, and compared this with the number of clusters obtained from Seurat using the default resolution; see Table 6 in Supplementary Note 2. For 4 out of the 8 RNA data sets evaluated in this article (RNAMix1, RNAMix2, BaronPanc, and CellMix), this procedure on PM_2_ yielded an estimate for *k* which matched the number of distinct annotated labels. On the other hand, Seurat correctly estimates the number of clusters for only 2 out of the 8 RNA data sets (RNAMix1 and TMLung).

Table 5 reports the geometric perturbation. We see that increasing *p* increases the geometric perturbation, with PM_1.5_ yielding the smallest geometric perturbation on all data sets. Although PM_1.5_ is the clear winner in terms of this metric, PM_2_ still performed favorably with respect to UMAP and *t*-SNE. Indeed, *r*d PM_2_ had lower geometric perturbation than UMAP on all but one data set (TMPanc), and lower geometric perturbation than *t*-SNE on the majority of data sets. Figure 4 shows the PCA, PM_2_, UMAP, and *t*-SNE embeddings of the Cell Mix data set, as well as a tree structure on the clusters. The tree structure was obtained by first computing the cluster means in the embedding and then applying hierarchical clustering with average linkage to the means. The PCA tree (Figure 4(e)) was computed using 40 PCs so that it accurately reflects the global geometry of the clusters. Interestingly path metrics recover the same hierarchical structure on the clusters as PCA: the cell types HCC827 and H1975 are the most similar, and H838 is the most distinct. This is what one would expect given more extensive biological information about the cell types, since H838 is the only cell line here derived from metastatic site Lymph node on a male patient, while both HCC827 and H1975 originated from the primary site of female lung cancer patients. However, neither UMAP or *t*-SNE give the correct hierarchical representation of the clusters, because both methods struggle to preserve global geometric structure as observed in numerous studies (83, 84). We note that in Figure 4(b) the clusters appear elongated in the PM_2_ embedding; such elongated cluster shapes occur when clusters living in nearly orthogonal subspaces (due for example to different genetic signatures) are projected into a lower-dimensional space; see Supplementary Note 3 for an example illustrating how this phenomenon occurs. While this is also the case for PCA, the PM embedding exaggerates the elongation by shrinking noisy directions. Although 2 dimensions is generally not sufficient to visualize the true cluster shapes, the PM embedding is able to simultaneously denoise the clusters while preserving their global layout.

Figure 5 records the runtime for processing and clustering (in minutes) of the Baron’s Pancreatic (*n* = 1844) and PBMC4K (*n* = 4316) data sets. For PBMC4k (our largest data set), we use the landmark-based approximation of path distances for scalability. All the PM methods run in less than a minute on BaronPanc and less than 6 minutes on PBMC4k; RaceID3, scanpy, and Seurat were also fast. SC3 and SIMLR had long runtimes, requiring 37.9 and 91.1 minutes respectively for PBMC4k.

**Fig. 5.**
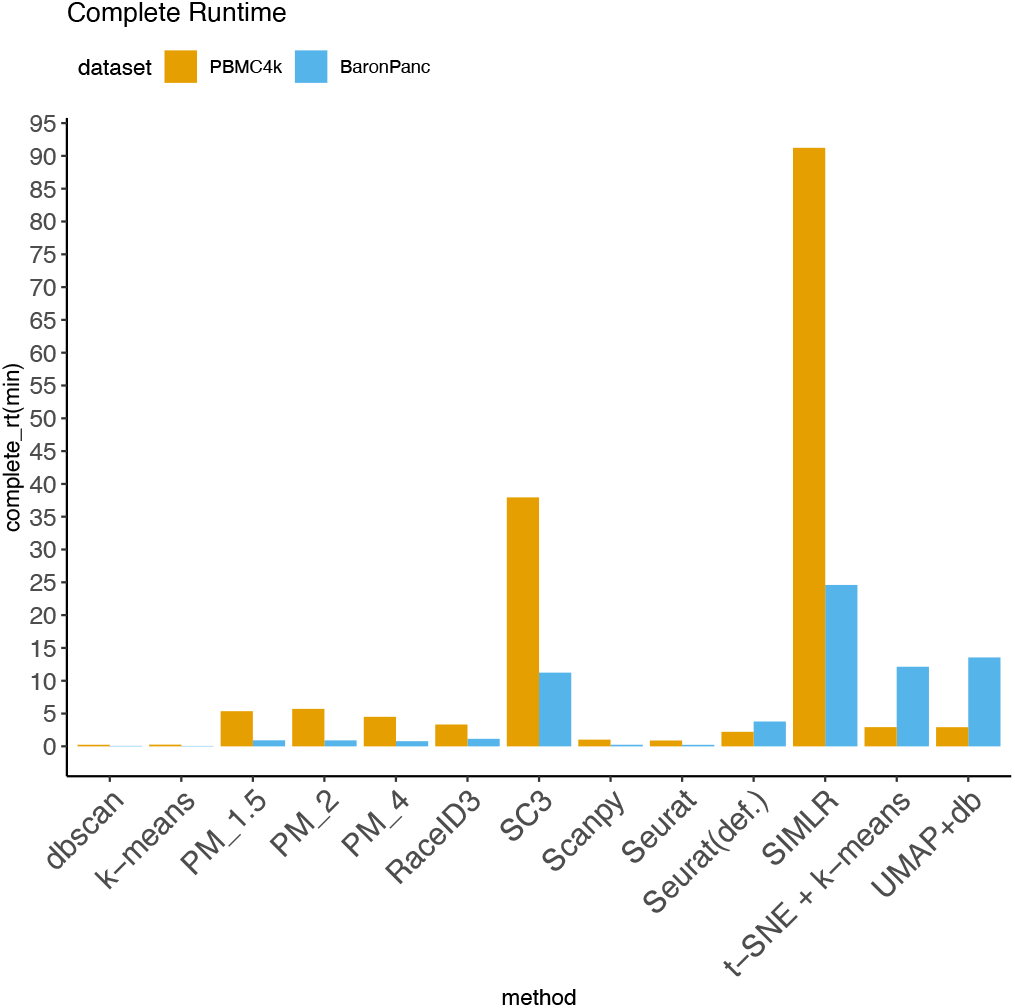
Processing and clustering time for PBMC4K and Baron’s Pancreatic data sets.

### Determining the parameter *p*

In this section, we explore the clustering performance of scPMP for different values of the parameter *p*. We record the ARI achieved by scPMP for each real data set for *p* ranging from 1 to 10 in increments of 0.5. Figure 6(a) plots the corresponding distributions of ARI; *p* =2 is the clear winner across various *p*, achieving the highest median ARI with the smallest spread of values. Furthermore, for each RNA data set we determined the *p* value maximizing the data set’s ARI and investigated whether there was a correlation between the best *p* and the degree of data elongation. We define an elongation score for each data set by computing the skewness coefficient of *k*th nearest neighbor distances for *k* = 10log(*n*). More specifically, letting 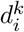 denote the Euclidean distance of *x*_*i*_ from its *k*th nearest neighbor, we define the data elongation score as the following measure of skewness:

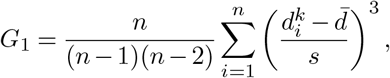

where 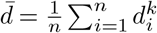 and *s* is the standard deviation of the 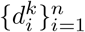.

**Fig. 6.**
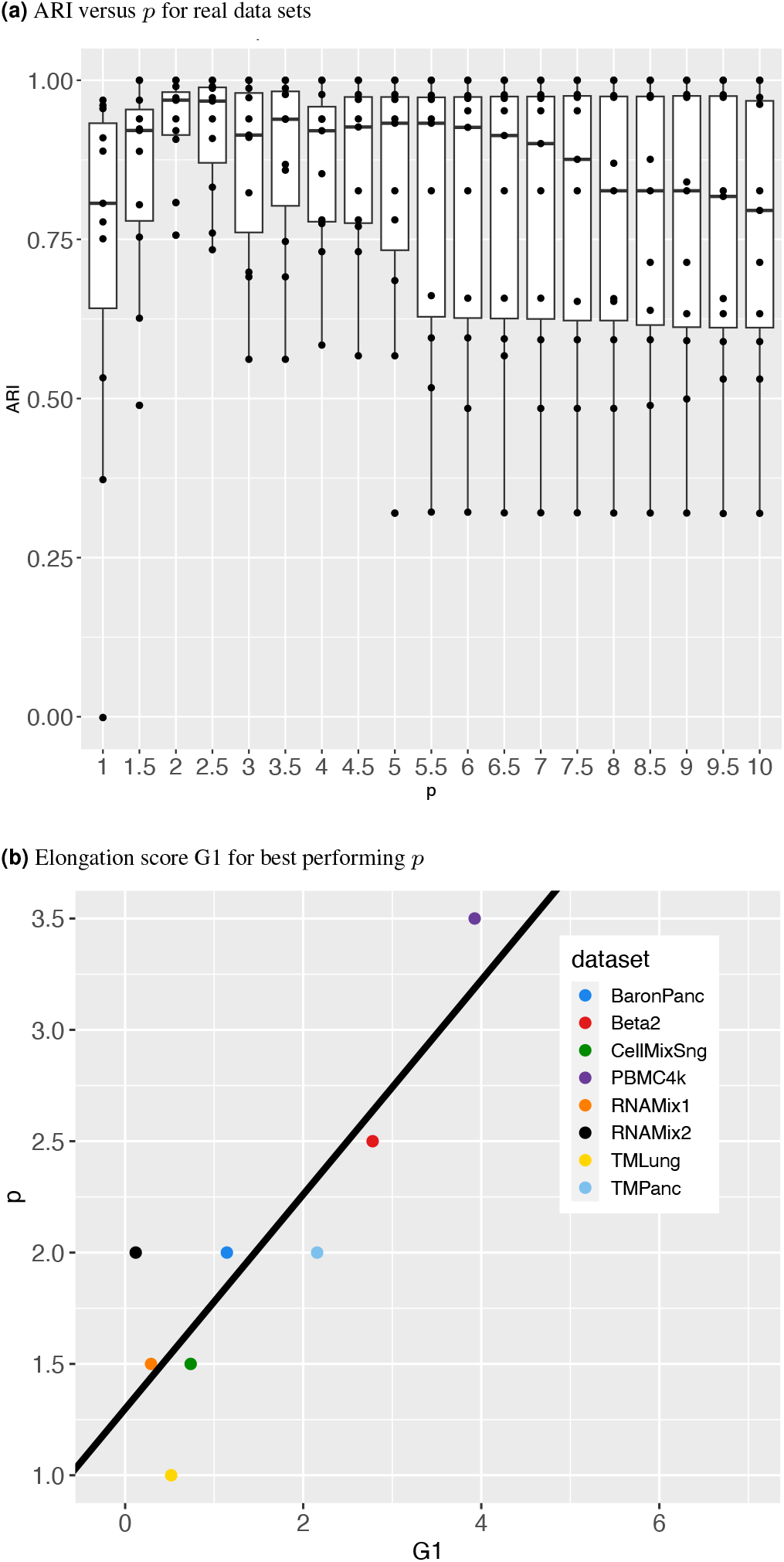
Clustering performance for different values of *p*

We observe a moderately strong linear relationship (*r* = 0.866) between the elongation score of a data set and the value of *p* achieving the best ARI as in Figure 6(b). Overall these results support using *p* =2 as a default, but increasing *p* if the data set exhibits strong elongation; the elongation score is a completely unsupervised statistic, and can thus be computed without access to data labels.

## Conclusions

This article introduces a new theoretical framework to analyze single-cell RNA-seq databased on the computation of optimal paths. Specifically, path metrics encode both geometric and density-based information, and the resulting low-dimensional embeddings simultaneously preserve density-based cluster structure as well as global cluster orientation. Thus, our method with theoretical guarantees addresses the inherent challenge of balancing the preservation of local cluster structures and the global data geometry, a common limitation in existing scRNAseq clustering and visualization methods such as DBSCAN, SC3, scanpy, and Seurat. The flexibility in choosing the parameter *p* allows researchers to adjust the balance between density sensitivity and geometry preservation, tailoring the analysis to their dataset’s specific characteristics, such as noise level and elongation. Compared to deep learning-based methods, such as CellPLM, scMoFormer, and scMoGNN, scPMP based on path metrics offers greater interpretability making it easier to derive biological insights. More importantly, scPMP is more robust to smaller datasets than deep learning-based methods since it has fewer parameters to be trained.

The method exhibits competitive performance when applied to numerous benchmarks, and the implementation is scalable to large data sets. Although we investigated other choices of *p*, we found that *p* = 2 performed well on a wide range of RNA data sets, indicating that *p* = 2 is an appropriate balance between density and geometry for this application.

Future research will explore ways to make the method more robust to noise, tools for better visualization of the PM embeddings, and adapting the method to the semi-supervised context.

## Software availability

The code to reproduce all reported results and generate figures is available at the scPMP github repository. The repository also contains a tutorial for the scPMP algorithm. Small differences observed during the reproduction of results is due to randomness introduced at the imputation step of data preprocessing.

## Data availability

The Cellmix and RNAmix data are downloaded from GEO under accession code GSE118767, and the preprocessed data are available at their github repository. The PBMC4K data is available at 10x Genomics’s website through https://support.10xgenomics.com/single-cell-gene-expression/datasets/2.1.0/pbmc4k. The Baron’s pancreatic data is available in GEO with the access code GSM2230757. The simulated data were created based on the Baron’s data. Simulation code is provided in the scPMP github repository. The mouse tissue scRNAseq data sets are accessible on Figshare.

## Bibliography

1. Antoine-Emmanuel Saliba, Alexander J. Westermann, Stanislaw A. Gorski, and Jörg Vogel. Single-cell RNA-seq: advances and future challenges. Nucleic Acids Research, 42(14): 8845–8860, 07 2014. ISSN 0305-1048. doi: 10.1093/nar/gku555.

2. James Eberwine, Hermes Yeh, Kevin Miyashiro, Yanxiang Cao, Suresh Nair, Richard Finnell, Martha Zettel, and Paul Coleman. Analysis of gene expression in single live neurons. Proceedings of the National Academy of Sciences, 89(7):3010–3014, 1992.

3. Fuchou Tang, Catalin Barbacioru, Yangzhou Wang, Ellen Nordman, Clarence Lee, Nanlan Xu, Xiaohui Wang, John Bodeau, Brian B Tuch, Asim Siddiqui, et al. mrna-seq whole-transcriptome analysis of a single cell. Nature methods, 6(5):377–382, 2009.

4. Vladimir Yu Kiselev, Kristina Kirschner, Michael T Schaub, Tallulah Andrews, Andrew Yiu, Tamir Chandra, Kedar N Natarajan, Wolf Reik, Mauricio Barahona, Anthony R Green, and Martin Hemberg. SC3: consensus clustering of single-cell RNA-seq data. Nature Methods, 14:483–486, 05 2017. doi: 10.1038/nmeth.4236.

5. Bo Wang, Junjie Zhu, Emma Pierson, Daniele Ramazzotti, and Serafim Batzoglou. Visualization and analysis of single-cell RNA-seq data by kernel-based similarity learning. Nature Methods, 14:414–416, 04 2017. doi: 10.1038/nmeth.4207.

6. Josip S Herman, Dominic Grün, et al. Fateid infers cell fate bias in multipotent progenitors from single-cell rna-seq data. Nature methods, 15(5):379, 2018.

7. Peijie Lin, Michael Troup, and Joshua W. K. Ho. CIDR: Ultrafast and accurate clustering through imputation for single-cell RNA-seq data. Genome Biology, 18, 03 2017. doi: 10.1186/s13059-017-1188-0.

8. Zeisel A, Ana B Muñoz-Manchado, Simone Codeluppi, Lönnerberg P, La Manno G, Juréus A, Marques S, Munguba H, He L, Betsholtz C, Rolny C, Castelo-Branco G, Hjerling-Leffler J, and Linnarsson S and. Brain structure. Cell types in the mouse cortex and hippocampus revealed by single-cell RNA-seq. Science (New York, N.Y.), 347:1138–1142, 03 2015. doi: 10.1126/science.aaa1934.

9. Justina žurauskieně and Christopher Yau. pcaReduce: hierarchical clustering of single cell transcriptional profiles. BMC Bioinformatics, 17, 03 2016. doi: 10.1186/s12859-016-0984-y.

10. Jacob H. CLevine, Erin F. Simonds, Sean C. Bendall, Kara L. Davis, El-ad D. Amir, Michelle D. Tadmor, Oren Litvin, Harris G. Fienberg, Astraea Jager, Eli R. Zunder, Rachel Finck, Amanda L. Gedman, Ina Radtke, James R. Downing, Dana Pe’er, and Garry P. Nolan. Data-driven phenotypic dissection of aml reveals progenitor-like cells that correlate with prognosis. Cell, 2015. doi: doi:10.1016/j.cell.2015.05.047.

11. Chen Xu and Zhengchang Su. Identification of cell types from single-cell transcriptomes using a novel clustering method. Bioinformatics, 31(12):1974–1980, 2015.

12. Tim Stuart, Andrew Butler, Paul Hoffman, Christoph Hafemeister, Efthymia Papalexi, William M Mauck III, Yuhan Hao, Marlon Stoeckius, Peter Smibert, and Rahul Satija. Comprehensive integration of single-cell data. Cell, 177:1888–1902, 2019. doi: 10.1016/j.cell.2019.05.031.

13. Xiaoshu Zhu, Jie Zhang, Yunpei Xu, Jianxin Wang, Xiaoqing Peng, and Hong-Dong Li. Single-cell clustering based on shared nearest neighbor and graph partitioning. Interdisciplinary Sciences: Computational Life Sciences, 12:117–130, 2020. doi: 10.1007/s12539-019-00357-4.

14. F. Alexander Wolf, Philipp Angerer, and Fabian J. Theis. SCANPY: large-scale single-cell gene expression data analysis. Genome Biology, 19, 02 2018. doi: 10.1186/s13059-017-1382-0.

15. Juexin Wang, Anjun Ma, Yuzhou Chang, Jianting Gong, Yuexu Jiang, Ren Qi, Cankun Wang, Hongjun Fu, Qin Ma, and Dong Xu. scgnn is a novel graph neural network framework for single-cell rna-seq analyses. Nature communications, 12(1):1882, 2021.

16. Romain Lopez, Jeffrey Regier, Michael B Cole, Michael I Jordan, and Nir Yosef. Deep generative modeling for single-cell transcriptomics. Nature methods, 15(12):1053–1058, 2018.

17. Tian Tian, Ji Wan, Qi Song, and Zhi Wei. Clustering single-cell rna-seq data with a model-based deep learning approach. Nature Machine Intelligence, 1(4):191–198, 2019.

18. Jiayuan Ding, Hongzhi Wen, Wenzhuo Tang, Renming Liu, Zhaoheng Li, Julian Venegas, Runze Su, Dylan Molho, Wei Jin, Wangyang Zuo, et al. Dance: A deep learning library and benchmark for single-cell analysis. bioRxiv, pages 2022–10, 2022.

19. Madalina Ciortan and Matthieu Defrance. GNN-based embedding for clustering scRNA-seq data. Bioinformatics, 38(4):1037–1044, 11 2021. ISSN 1367-4803. doi: 10.1093/bioinformatics/btab787.

20. Yuansong Zeng, Xiang Zhou, Jiahua Rao, Yutong Lu, and Yuedong Yang. Accurately clustering single-cell rna-seq data by capturing structural relations between cells through graph convolutional network. In 2020 IEEE International Conference on Bioinformatics and Biomedicine (BIBM), pages 519–522, 2020. doi: 10.1109/BIBM49941.2020.9313569.

21. Tian Tian, Jie Zhang, Xiang Lin, Zhi Wei, and Hakon Hakonarson. Model-based deep embedding for constrained clustering analysis of single cell rna-seq data. Nature communications, 12(1):1873, 2021.

22. Xiangjie Li, Kui Wang, Yafei Lyu, Huize Pan, Jingxiao Zhang, Dwight Stambolian, Katalin Susztak, Muredach P Reilly, Gang Hu, and Mingyao Li. Deep learning enables accurate clustering with batch effect removal in single-cell rna-seq analysis. Nature communications, 11(1):2338, 2020.

23. Duc Tran, Hung Nguyen, Bang Tran, Carlo La Vecchia, Hung N Luu, and Tin Nguyen. Fast and precise single-cell data analysis using a hierarchical autoencoder. Nature communications, 12(1):1029, 2021.

24. Liang Chen, Weinan Wang, Yuyao Zhai, and Minghua Deng. Deep soft k-means clustering with self-training for single-cell rna sequence data. NAR genomics and bioinformatics, 2(2): lqaa039, 2020.

25. Yanglan Gan, Xingyu Huang, Guobing Zou, Shuigeng Zhou, and Jihong Guan. Deep structural clustering for single-cell rna-seq data jointly through autoencoder and graph neural network. Briefings in Bioinformatics, 23(2):bbac018, 2022.

26. Hongzhi Wen, Wenzhuo Tang, Xinnan Dai, Jiayuan Ding, Wei Jin, Yuying Xie, and Jiliang Tang. Cellplm: Pre-training of cell language model beyond single cells. bioRxiv, pages 2023–10, 2023.

27. Wenzhuo Tang, Renming Liu, Hongzhi Wen, Xinnan Dai, Jiayuan Ding, Hang Li, Wenqi Fan, Yuying Xie, and Jiliang Tang. A general single-cell analysis framework via conditional diffusion generative models. bioRxiv, pages 2023–10, 2023.

28. Hongzhi Wen, Jiayuan Ding, Wei Jin, Yiqi Wang, Yuying Xie, and Jiliang Tang. Graph neural networks for multimodal single-cell data integration. In Proceedings of the 28th ACM SIGKDD conference on knowledge discovery and data mining, pages 4153–4163, 2022.

29. Wenzhuo Tang, Hongzhi Wen, Renming Liu, Jiayuan Ding, Wei Jin, Yuying Xie, Hui Liu, and Jiliang Tang. Single-cell multimodal prediction via transformers. arXiv preprint 2303.00233, 2023.

30. Zhuohan Yu, Yifu Lu, Yunhe Wang, Fan Tang, Ka-Chun Wong, and Xiangtao Li. Zinb-based graph embedding autoencoder for single-cell rna-seq interpretations. In Proceedings of the AAAI conference on artificial intelligence, pages 4671–4679, 2022.

31. Dylan Molho, Jiayuan Ding, Wenzhuo Tang, Zhaoheng Li, Hongzhi Wen, Yixin Wang, Julian Venegas, Wei Jin, Renming Liu, Runze Su, et al. Deep learning in single-cell analysis. ACM Transactions on Intelligent Systems and Technology, 2022.

32. Leland McInnes, John Healy, and James Melville. Umap: Uniform manifold approximation and projection for dimension reduction. arXiv preprint 1802.03426, 2018.

33. Laurens Van der Maaten and Geoffrey Hinton. Visualizing data using t-sne. Journal of machine learning research, 9(11), 2008.

34. Kevin R. Moon, David van Dijk, Zheng Wang, Scott Gigante, Daniel B. Burkhardt, William S. Chen, Kristina Yim, Antonia van den Elzen, Matthew J. Hirn, Ronald R. Coifman, Natalia B. Ivanova, Guy Wolf, and Smita Krishnaswamy. Visualizing structure and transitions in high-dimensional biological data. Nature Biotechnology, 37(12):1482–1492, 2019. doi: 10.1038/s41587-019-0336-3.

35. Jian Tang, Jingzhou Liu, Ming Zhang, and Qiaozhu Mei. Visualizing large-scale and high-dimensional data. In Proceedings of the 25th international conference on world wide web, pages 287–297, 2016.

36. Luyi Tian, Xueyi Dong, Saskia Freytag, Kim-Anh Lê Cao, Shian Su, Abolfazl JalalAbadi, Daniela Amann-Zalcenstein, Tom S Weber, Azadeh Seidi, Jafar S Jabbari, et al. Bench-marking single cell rna-sequencing analysis pipelines using mixture control experiments. Nature methods, 16(6):479–487, 2019.

37. Martin Ester, Hans-Peter Kriegel, Jörg Sander, Xiaowei Xu, et al. A density-based algorithm for discovering clusters in large spatial databases with noise. In Kdd, volume 96, pages 226–231, 1996.

38. Xiaowei Xu, Martin Ester, H-P Kriegel, and Jörg Sander. A distribution-based clustering algorithm for mining in large spatial databases. In Proceedings 14th International Conference on Data Engineering, pages 324–331. IEEE, 1998.

39. Logistical coordination et al. Tabula Muris Consortium, Overall coordination. Single-cell transcriptomics of 20 mouse organs creates a tabula muris. Nature, 562:367–372, 2018.

40. P. Vincent and Y. Bengio. Density-sensitive metrics and kernels. In Snowbird Learning Workshop, 2003.

41. O. Bousquet, O. Chapelle, and M. Hein. Measure based regularization. In NIPS, pages 1221–1228, 2004.

42. Sajama and A. Orlitsky. Estimating and computing density based distance metrics. In ICML, pages 760–767, 2005.

43. H. Chang and D.-Y. Yeung. Robust path-based spectral clustering. Pattern Recognition, 41 (1):191–203, 2008.

44. A.S. Bijral, N. Ratliff, and N. Srebro. Semi-supervised learning with density based distances. In UAI, pages 43–50, 2011.

45. A. Moscovich, A. Jaffe, and B. Nadler. Minimax-optimal semi-supervised regression on unknown manifolds. In AISTATS, pages 933–942, 2017.

46. D. Mckenzie and S. Damelin. Power weighted shortest paths for clustering Euclidean data. Foundations of Data Science, 1(3):307, 2019.

47. A. Little, M. Maggioni, and J.M Murphy. Path-based spectral clustering: Guarantees, robustness to outliers, and fast algorithms. Journal of Machine Learning Research, 21(6):1–66, 2020.

48. Ximena Fernández, Eugenio Borghini, Gabriel Mindlin, and Pablo Groisman. Intrinsic persistent homology via density-based metric learning. Journal of Machine Learning Research, 24(75):1–42, 2023.

49. Bernd Fischer, Thomas Zöller, and Joachim M Buhmann. Path based pairwise data clustering with application to texture segmentation. In International Workshop on Energy Minimization Methods in Computer Vision and Pattern Recognition, pages 235–250. Springer, 2001.

50. S. Zhang and J.M. Murphy. Hyperspectral image clustering with spatially-regularized ultrametrics. Remote Sensing, 13(5):955, 2021.

51. M. Pollack. Letter to the editor: The maximum capacity through a network. Operations Research, 8(5):733–736, 1960.

52. T.C. Hu. Letter to the editor: The maximum capacity route problem. Operations Research, 9(6):898–900, 1961.

53. P.M. Camerini. The min-max spanning tree problem and some extensions. Information Processing Letters, 1(10-14), 1978.

54. H. Gabow and R.E. Tarjan. Algorithms for two bottleneck optimization problems. Journal of Algorithms, 9:411–417, 1988.

55. J.B. Tenenbaum, V. De Silva, and J.C. Langford. A global geometric framework for nonlinear dimensionality reduction. Science, 290(5500):2319–2323, 2000.

56. John M Lee. Introduction to Riemannian manifolds. Springer, 2018.

57. S.J. Hwang, S.B. Damelin, and A. Hero. Shortest path through random points. The Annals of Applied Probability, 26(5):2791–2823, 2016.

58. Pablo Groisman, Matthieu Jonckheere, and Facundo Sapienza. Nonhomogeneous euclidean first-passage percolation and distance learning. Bernoulli, 28(1):255–276, 2022.

59. Ximena Fernández, Eugenio Borghini, Gabriel Mindlin, and Pablo Groisman. Intrinsic persistent homology via density-based metric learning. Journal of Machine Learning Research, 24(75):1–42, 2023.

60. Timothy Chu, Gary Miller, and Donald Sheehy. Exploration of a graph-based density sensitive metric. arXiv preprint 1709.07797, 2017.

61. Anna Little, Daniel McKenzie, and James M Murphy. Balancing geometry and density: Path distances on high-dimensional data. SIAM Journal on Mathematics of Data Science, 4(1): 72–99, 2022.

62. Pablo Groisman, Matthieu Jonckheere, and Facundo Sapienza. Nonhomogeneous euclidean first-passage percolation and distance learning. Bernoulli, 28(1):255–276, 2022.

63. T. Chu, G.L. Miller, and D.R. Sheehy. Exact computation of a manifold metric, via Lipschitz embeddings and shortest paths on a graph. In SODA, pages 411–425, 2020.

64. Moshe Sniedovich. Dijkstra’s algorithm revisited: the dynamic programming connexion. Control and cybernetics, 35(3):599–620, 2006.

65. N. García Trillos, D. Sanz-Alonso, and R. Yang. Local regularization of noisy point clouds: Improved global geometric estimates and data analysis. Journal of Machine Learning Research, 20(136):1–37, 2019.

66. Benyamin Ghojogh, Ali Ghodsi, Fakhri Karray, and Mark Crowley. Multidimensional scaling, sammon mapping, and isomap: Tutorial and survey. 2020.

67. Clifford Lam and Qiwei Yao. Factor modeling for high-dimensional time series: inference for the number of factors. The Annals of Statistics, pages 694–726, 2012.

68. Christopher Williams and Matthias Seeger. Using the nyström method to speed up kernel machines. In Proceedings of the 14th annual conference on neural information processing systems, number CONF, pages 682–688, 2001.

69. Benyamin Ghojogh, Ali Ghodsi, Fakhri Karray, and Mark Crowley. Multidimensional scaling, sammon mapping, and isomap: Tutorial and survey. arXiv preprint 2009.08136, 2020.

70. John Platt. Fastmap, metricmap, and landmark mds are all nyström algorithms. In International Workshop on Artificial Intelligence and Statistics, pages 261–268. PMLR, 2005.

71. H Yu, X Zhao, X Zhang, and Y Yang. Isomap using nyström method with incremental sampling. Advances in Information Sciences & Service Sciences, 4(12), 2012.

72. Ali Civril, Malik Magdon-Ismail, and Eli Bocek-Rivele. Ssde: Fast graph drawing using sampled spectral distance embedding. In International Symposium on Graph Drawing, pages 30–41. Springer, 2006.

73. Gil Shamai, Michael Zibulevsky, and Ron Kimmel. Efficient inter-geodesic distance computation and fast classical scaling. IEEE Transactions on Pattern Analysis and Machine Intelligence, 42(1):74–85, 2020. doi: 10.1109/TPAMI.2018.2877961.

74. Leonard Kaufman and Peter Rousseeuw. Finding Groups in Data: An Introduction to Cluster Analysis. 09 2009. ISBN 9780470317488.

75. Martin Maechler, Peter Rousseeuw, Anja Struyf, Mia Hubert, and Kurt Hornik. cluster: Cluster Analysis Basics and Extensions, 2021.

76. Ulrike Von Luxburg. A tutorial on spectral clustering. Statistics and computing, 17(4):395–416, 2007.

77. Dominic Grün et al. Revealing dynamics of gene expression variability in cell state space. Nature methods, 17:45–49, 2018.

78. Mo Huang, Jingshu Wang, Eduardo Torre, Hannah Dueck, Sydney Shaffer, Roberto Bonasio, John I Murray, Arjun Raj, Mingyao Li, and Nancy R Zhang. Saver: gene expression recovery for single-cell rna sequencing. Nature methods, 15(7):539–542, 2018.

79. Wei Vivian Li and Jingyi Jessica Li. An accurate and robust imputation method scimpute for single-cell rna-seq data. Nature communications, 9(1):1–9, 2018.

80. Maayan Baron et al. A single-cell transcriptomic map of the human and mouse pancreas reveals inter-and intra-cell population structure. Cell Systems, 3(4):346–360, 2016.

81. Christoph Hafemeister and Rahul Satija. Normalization and variance stabilization of single-cell rna-seq data using regularized negative binomial regression. Genome Biology, 20(1), 2019.

82. Saket Choudhary and Rahul Satija. Comparison and evaluation of statistical error models for scrna-seq. Genome Biology, 23, 2022.

83. Dmitry Kobak and Philipp Berens. The art of using t-sne for single-cell transcriptomics. Nature Communications, 10:2041–1723, 2019. doi: 10.1038/s41467-019-13056-x.

84. Shamus M Cooley, Timothy Hamilton, Samuel D Aragones, J Christian J Ray, and Eric J Deeds. A novel metric reveals previously unrecognized distortion in dimensionality reduction of scrna-seq data. Biorxiv, page 689851, 2019.

