## Supplemental Material for "Clustering and visualization of single-cell RNA-seq data using path metrics"

### Supplementary Note 1: Data Pre-processing

In this section the pre-processing of all RNA data sets is described. The main preprocessing steps are quality control, imputation with SAVER (1), and normalization. Below we provide information about quality control and imputation and then we describe how we used those steps according to the guidelines of each method.

**Main steps. Quality Control:** Quality control is applied on RNAmix1, RNAmix2, Cellmix, BaronPanc, PMC4K, Beta. Specifically, cells where at most 200 genes are expressed are filtered out. Also, only genes that are expressed in more than 3 cells are included in the data set. In addition, cells with percentage of expressed mitochondrial genes greater than 20% are excluded. The data sets TMpanc and TMLung as found in Figshare have passed a quality control check with cutoffs of at least 500 genes and 50,000 reads, so no additional filtering was applied.

**Imputation:** Imputation with SAVER (1) was applied to all RNA seq data sets apart from Cellmix. After removing multiplets the Cellmix data set included high quality data and every clustering method achieved high ARI, suggesting no need for further processing and imputation.

**Preprocessing per method. Path metrics (PM), $k$-means, DBSCAN**: After quality control and imputation, we normalize the data. RNAmix1, RNAmix2, TMLung, Beta, TMPanc, PBMC4K were row normalized and log transformed (data matrix had cells in rows and genes in columns). We then restrict to the top 2000 high variance genes. For the BaronPanc and CellMix, which have large sample size, SCT transformation was applied instead and the top 3000 variable genes were kept (2, 3). When needed, we rescale genes where variances were extremely high. As a next step we apply PCA for dimension reduction, keeping the top 40 PC's. Finally, denoising is applied by replacing each point with the mean of its local neighborhood, using a neighborhood size of $K = 12$ points. For very large data sets, one may want to use a larger $K$.

**UMAP+DBSCAN, $t$-SNE+$k$-means**: After quality control and imputation, we apply Linnnorm (4) to all data sets. Then, we restrict to the top 2000 high variance genes. When needed, we rescale genes with extremely high variance. Finally, we apply PCA for dimension reduction, keeping the top 40 PC's.

**Seurat**: For this method, we process the data as for PM and then use Seurat's(5) functions to find neighboring points and cluster them. Notice that here we adjust the parameter 'res', to retrieve the correct number of clusters.

**Seurat_def**: We follow the suggested processing and clustering workflow of Seurat (5) for all data sets. Notice that we normalize BaronPanc and CellMix with the SCT method (2, 3). Then data sets are clustered with adjusted resolution parameter, to retrieve the correct number of clusters.

**SC3**: After quality control and imputation we normalize the information of every cell and multiply by 10000. Then we use the log of the data for clustering with SC3 (6). Exception to this are the BaronPanc and CellMix data set, for which we use SCT normalization.

**scanpy**: After quality control and imputation we use the lognormalization of scanpy (7). Exception to this are the BaronPanc and CellMix data set, for which we use SCT normalization.

**RaceID3**: We apply quality control on the cells of the counts of the data set. RaceID3 (8, 9) applies filtering and normalization in one step, which we adjust to have about the same amount of cells and genes as with other methods. Notice that we do not apply imputation because imputed data would not be counts, which are the required input of RaceID3.

**SIMLR**: For SIMLR (10) After quality control and imputation we normalize the information of every cell and multiply by 10000 and use the logorithm of those data. Exception to this are the BaronPanc and CellMix data set, for which we use SCT normalization.

#### Denoising with local averaging.

As a denoising step, the scPMP algorithm performs local averaging. Specifically, the observed feature of a point is substituted for the average of that feature across the $K_1$ nearest neighbors of the point. The effect of local averaging on clustering with scPMP was explored, and it was found that local averaging not only contributes to an increase in ARI, but it also reduces the overall runtime of the algorithm (Table 1). Furthermore, we explore the runtime and clustering performance of scPMP when coupled with other preprocessing methods than those mentioned in appendix 1. In more detail, we use the normalization and dimension reduction process of scVI (11), which assumes that expression data are generated from a Zero Inflated Negative Binomial (ZINB) model and estimates the parameters of this model by applying variational inference. We additionally investigate the effect of scVI preprocessing on scPMP clustering with and without local averaging. According to our results in Table 2, scVI yielded better ARI values when combined with local averaging for most cases (Baron(p=2), PBMC4K(p=2), and didn't outperform scPMP clustering with the default preprocessing and denoising methods (as seen in the last two rows of Table 1).

**Table 1.** Clustering performance with and without local averaging denoising

| Dataset | Local Averaging | ARI | ECP | ECA | complete runtime | clustering runtime |
|---|---|---|---|---|---|---|
| BaronPanc | false | 0.791 | 0.289 | 0.158 | 4.643 | 4.197 |
| PBMC4k | false | 0.744 | 0.321 | 0.196 | 18.077 | 12.844 |
| BaronPanc | true | **0.969** | 0.081 | 0.077 | 0.739 | **0.682** |
| PBMC4k | true | **0.978** | 0.055 | 0.096 | 4.936 | **4.702** |

**Table 2.** Clustering performance after preprocessing with scVI

| Dataset | Local Averaging | ARI | ECP | ECA | complete runtime | clustering runtime |
|---|---|---|---|---|---|---|
| BaronPanc(p=2) | false | 0.653 | 0.440 | 0.332 | 2.436 | 2.300 |
| BaronPanc(p=2) | true | 0.721 | 0.373 | 0.277 | 2.386 | 2.217 |
| PBMC4k(p=4) | false | 0.745 | 0.324 | 0.201 | 12.550 | 12.132 |
| PBMC4k(p=4) | true | 0.741 | 0.320 | 0.188 | 11.109 | 10.745 |
| PBMC4k(p=2) | false | 0.744 | 0.319 | 0.191 | 13.475 | 13.072 |
| PBMC4k(p=2) | true | 0.748 | 0.315 | 0.190 | 12.732 | 12.348 |

### Supplementary Note 2: Additional Clustering Results

Here we present more clustering evaluation results based on Entropy of Cluster Accuracy (ECA) and Entropy of cluster Purity (ECP). The ECA can quantify the variety of true labels within a predicted cluster and ECP can quantify the variety of predicted cluster labels within a true group.

*Definition 1:* Let $N$ represent the number of true groups and $M$ the number of predicted clusters. Let $N_j$ be the number of true groups with data points within the $j^{\text{th}}$ predicted cluster and similarly let $M_j$ be the number of predicted clusters with data points within the $j^{\text{th}}$ true group. Finally let $p(x_j)$ denote the proportion of data points belonging to the $j^{\text{th}}$ true group that are within a given $j^{\text{th}}$ predicted cluster and let $p_i(y_j)$ denote the proportion of data points of $j^{\text{th}}$ predicted cluster that are within a given $i^{\text{th}}$ true group. Then:

$$\text{ECA} = -\frac{1}{M}\sum_{i=1}^{M}\sum_{j=1}^{N_i} p_i(x_j)\log(p(x_j)),$$

$$\text{ECP} = -\frac{1}{N}\sum_{i=1}^{N}\sum_{j=1}^{M_i} p_i(y_j)\log(p(y_j)).$$

For a given clustering, low ECA means that data points in a predicted cluster originate from the same true group. On the other hand, low ECP indicates that almost all the data points in a true group were assigned the same clustering label. Use of ECP and ECA in clustering of scRNAseq data was also found in (12).

**Table 3.** ECP for manifold data

| Method | Balls | EWB | Swiss | SO(3) |
|---|---|---|---|---|
| $k$-means | **0.082** | 1.050 | 0.588 | 1.084 |
| DBSCAN | 0.385 | 0.114 | **0** | **0** |
| UMAP+DBSCAN | 0.941 | 0.695 | **0** | **0** |
| $t$-SNE+ $k$-means | 0.153 | 0.630 | **0** | 0.440 |
| Seurat | 0.255 | 0.193 | **0** | **0** |
| $\text{PM}_{1.5}$ | 0.123 | 0.447 | **0** | 0.460 |
| $\text{PM}_2$ | 0.142 | **0.020** | **0** | **0** |
| $\text{PM}_4$ | 0.253 | 0.268 | **0** | **0** |

**Table 4.** ECA for manifold data

| Method | Balls | EWB | Swiss | SO(3) |
|---|---|---|---|---|
| $k$-means | **0.082** | 1.096 | 0.633 | 1.089 |
| DBSCAN | 0.362 | 0.231 | **0** | **0** |
| UMAP+DBSCAN | 0.200 | **0.014** | **0** | **0** |
| $t$-SNE+ $k$-means | 0.147 | 0.582 | **0** | 0.440 |
| Seurat | 0.250 | 0.183 | **0** | **0** |
| $\text{PM}_{1.5}$ | 0.120 | 0.461 | **0** | 0.462 |
| $\text{PM}_2$ | 0.138 | 0.020 | **0** | **0** |
| $\text{PM}_4$ | 0.248 | 0.291 | **0** | **0** |

**Table 5.** ECP for RNA data

| Method | RNA1 | RNA2 | TMLung | Beta | TMPanc | BaronPanc | PBMC4k | CellMix |
|---|---|---|---|---|---|---|---|---|
| SC3 | 0.328 | 0.114 | 0.294 | 0.058 | 0.070 | 0.301 | 0.062 | 0 |
| Scanpy | 0.517 | 0.183 | 0.322 | 0.128 | 0.516 | 0.088 | 0.057 | 0 |
| RaceID3 | 0.381 | 0.665 | **0.182** | 0.351 | 0.268 | 0.413 | 0.310 | 0 |
| SIMLR | 0.151 | 0.267 | 0.275 | 0.048 | 0.543 | 0.380 | 0.360 | 0 |
| Seurat | 0.292 | 0.282 | 0.230 | 0.155 | 0.540 | 0.122 | 0.053 | 0.027 |
| Seurat_def | 0.320 | 0.258 | 0.436 | 0.114 | 0.284 | 0.089 | 0.062 | 0 |
| $k$-means | 0.131 | 0.255 | 0.244 | 0.058 | 0.215 | 0.395 | 0.316 | 0 |
| DBSCAN | **0.075** | 0.141 | 0.404 | 0.083 | 0.138 | 0.109 | **0.051** | 0 |
| UMAP+db | 0.151 | 0.226 | 0.413 | **0.023** | **0.061** | 0.248 | 0.087 | 0 |
| $t$-SNE+ $k$-means | 0.102 | 0.133 | 0.437 | 0.052 | 0.494 | 0.402 | 0.451 | 0.147 |
| $\text{PM}_{1.5}$ | 0.096 | 0.136 | 0.197 | 0.058 | 0.482 | 0.273 | 0.312 | 0 |
| $\text{PM}_2$ | 0.096 | **0.062** | 0.323 | 0.058 | 0.141 | **0.081** | 0.308 | **0** |
| $\text{PM}_4$ | 0.096 | 0.114 | 0.184 | 0.108 | 0.260 | 0.226 | 0.055 | 0 |

**Table 6.** Predicted number of clusters for Seurat and Path metrics for RNA data ($k$ is the true number of clusters).

| Method | RNA1 | RNA2 | TMLung | Beta | TMPanc | BaronPanc | PBMC4k | CellMix |
|---|---|---|---|---|---|---|---|---|
| Seurat_res=0.8 | **7** | 8 | **7** | 6 | 11 | 12 | 13 | 14 |
| $\text{PM}_{1.5}$ | 12 | 11 | 8 | 4 | 15 | 9 | **4** | **5** |
| $\text{PM}_2$ | **7** | **7** | 9 | 4 | 5 | **8** | 5 | **5** |
| $\text{PM}_4$ | 8 | 8 | 16 | 4 | 5 | 7 | **4** | **5** |
| True $k$ | 7 | 7 | 7 | 3 | 7 | 8 | 4 | 5 |

**Table 7.** ECA for RNA data

| Method | RNA1 | RNA2 | TMLung | Beta | TMPanc | BaronPanc | PBMC4k | CellMix |
|---|---|---|---|---|---|---|---|---|
| SC3 | 0.289 | 0.114 | 0.228 | 0.058 | 0.132 | 0.368 | 0.328 | 0 |
| Scanpy | 0.481 | 0.242 | 0.314 | 0.129 | 0.309 | 0.093 | **0.054** | 0 |
| RaceID3 | 0.336 | 0.621 | **0.168** | 0.342 | **0.122** | 0.181 | 0.207 | 0.000 |
| SIMLR | 0.163 | 0.294 | 0.263 | 0.049 | 0.407 | 0.104 | 0.190 | 0 |
| Seurat | 0.319 | 0.230 | 0.193 | 0.153 | 0.290 | 0.097 | 0.265 | 0 |
| Seurat_def | 0.256 | 0.270 | 0.423 | 0.109 | 0.289 | 0.112 | 0.106 | 0 |
| $k$-means | 0.147 | 0.268 | 0.221 | 0.058 | 0.194 | 0.164 | 0.193 | 0 |
| DBSCAN | 0.090 | 0.188 | 0.368 | 0.465 | 0.202 | 0.146 | 0.262 | 0 |
| UMAP+db | **0.078** | 0.151 | 0.449 | 0.364 | 0.124 | **0.076** | 0.163 | 0 |
| $t$-SNE+ $k$-means | 0.104 | 0.137 | 0.426 | **0.052** | 0.259 | 0.171 | 0.187 | 0.126 |
| $PM_{1.5}$ | 0.110 | 0.146 | 0.180 | 0.058 | 0.305 | 0.147 | 0.196 | 0 |
| $PM_2$ | 0.110 | **0.071** | 0.362 | 0.058 | 0.196 | 0.077 | 0.195 | **0** |
| $PM_4$ | 0.110 | 0.123 | 0.230 | 0.106 | 0.156 | 0.159 | 0.096 | 0 |

**Table 8.** Downsampling results

| Dataset | Seurat | Seurat_def | $PM_{1.5}$ | $PM_2$ | $PM_4$ |
|---|---|---|---|---|---|
| 100% of Baron's Pancreatic | 0.941 | 0.971 | 0.804 | 0.969 | 0.853 |
| 50% of Baron's Pancreatic | 0.880 | 0.844 | 0.969 | 0.969 | 0.969 |
| 25% of Baron's Pancreatic | 0.973 | 0.705 | 0.973 | 0.973 | 0.973 |
| 10% of Baron's Pancreatic | 0.410 | 0.185 | 0.674 | 0.939* | 0.804 |

### Supplementary Note 3: Clustering visualizations on PCA and scPMP embedding

Figure 2 showes the scPMP embeddings of our benchmarking data sets colored by both the ground truth and predicted labels. For many of the data sets, some clusters appear as elongated bars in the $PM_2$ embedding. This tends to occur when clusters live in nearly orthogonal spaces due to different genetic signatures, i.e. they have high/low expression values on disjoint sets of genes. This effect also occurs for PCA, but the denoising effect of the $PM_2$ embedding exaggerates the effect. Consider for example 3 clusters, where $C_1$ expresses high values of $x_1, x_2$, $C_2$ expresses high values of $x_3, x_4$, and $C_3$ expresses high values of $x_5, x_6$, but all three clusters have "noise" appearing in all dimensions. Figure 1 shows the corresponding PCA and $PM_2$ plots (uniform sampling was performed in each coordinate). Note the PCA plot is elongated due to the approximate orthogonality, but the PM embedding exaggerates this effect by stretching the data in regions of high density (i.e. "useful" directions), and shrinking in noisy directions where the data is sparse. Thus the clusters in the $PM_2$ plot appear even more elongated. However the bar like appearance is not indicative of the true shape of the clusters but an artifact of the projection into the lower-dimensional space.

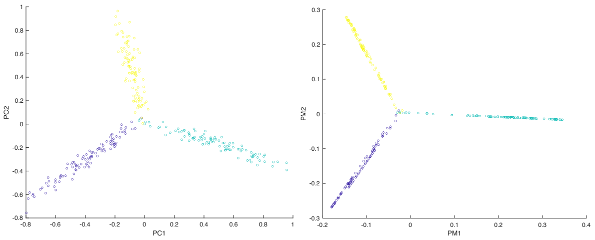


**Fig. 1.** Elongation effect for PCA and $PM_2$ embeddings

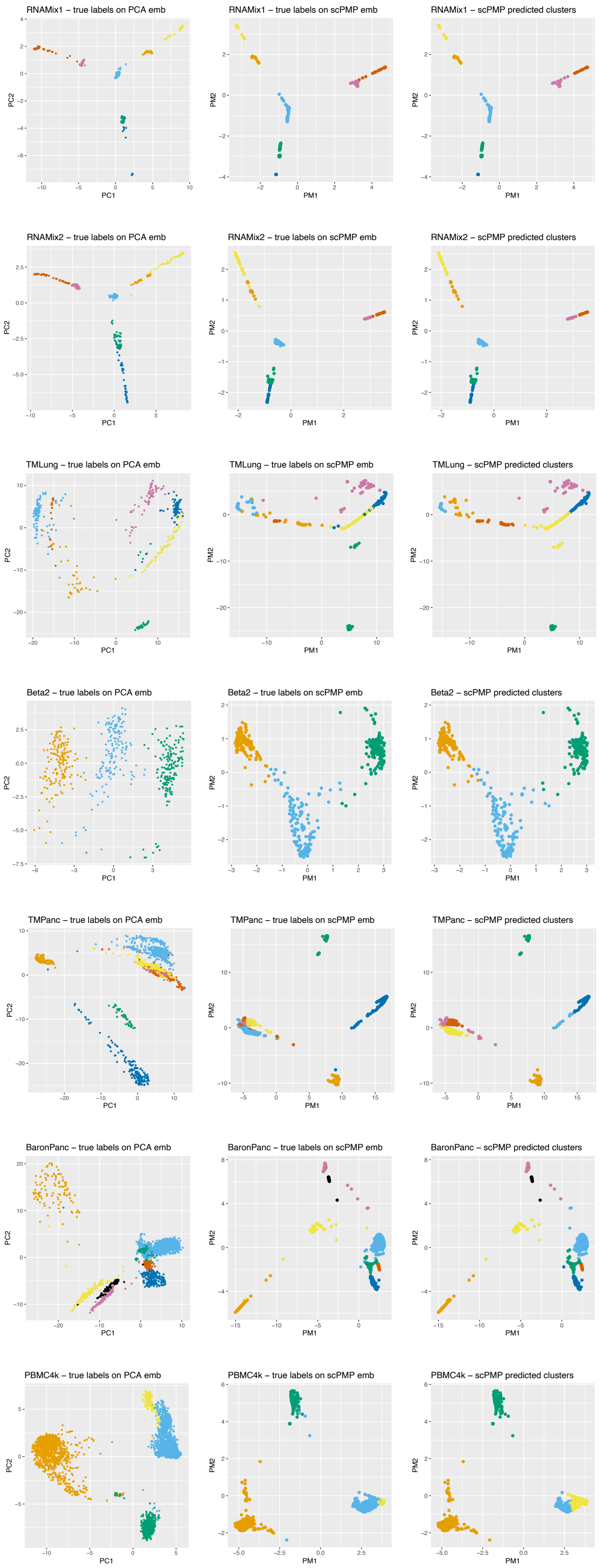


**Fig. 2.** Clustering results on PCA and scPMP embedding for real data

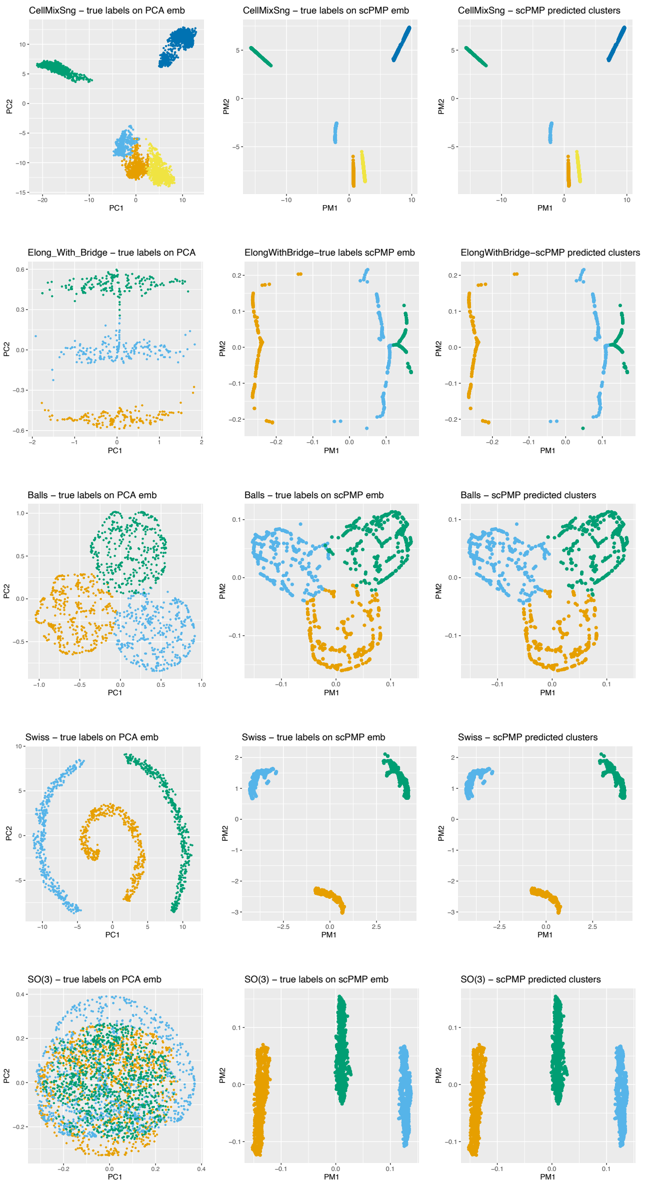


**Fig. 2.** Clustering results on PCA and scPMP embedding (continuing)


1. Mo Huang, Jingshu Wang, Eduardo Torre, Hannah Dueck, Sydney Shaffer, Roberto Bonasio, John I Murray, Arjun Raj, Mingyao Li, and Nancy R Zhang. Saver: gene expression recovery for single-cell rna sequencing. *Nature methods*, 15(7):539–542, 2018.
2. Christoph Hafemeister and Rahul Satija. Normalization and variance stabilization of single-cell rna-seq data using regularized negative binomial regression. *Genome Biology*, 20(1), 2019.
3. Saket Choudhary and Rahul Satija. Comparison and evaluation of statistical error models for scrna-seq. *Genome Biology*, 23, 2022.
4. Shun H. Yip, Panwen Wang, Jean-Pierre A. Kocher, Pak Chung Sham, and Junwen Wang. Linnorm: improved statistical analysis for single cell RNA-seq expression data. *Nucleic Acids Research*, 45(22):e179–e179, 09 2017. ISSN 0305-1048. doi: 10.1093/nar/gkx828.
5. Tim Stuart, Andrew Butler, Paul Hoffman, Christoph Hafemeister, Efthymia Papalexi, William M Mauck III, Yuhan Hao, Marlon Stoeckius, Peter Smibert, and Rahul Satija. Comprehensive integration of single-cell data. *Cell*, 177:1888–1902, 2019. doi: 10.1016/j.cell.2019.05.031.
6. Vladimir Yu Kiselev, Kristina Kirschner, Michael T Schaub, Tallulah Andrews, Andrew Yiu, Tamir Chandra, Kedar N Natarajan, Wolf Reik, Mauricio Barahona, Anthony R Green, and Martin Hemberg. SC3: consensus clustering of single-cell RNA-seq data. *Nature Methods*, 14:483–486, 05 2017. doi: 10.1038/nmeth.4236.
7. F. Alexander Wolf, Philipp Angerer, and Fabian J. Theis. SCANPY: large-scale single-cell gene expression data analysis. *Genome Biology*, 19, 02 2018. doi: 10.1186/s13059-017-1382-0.
8. Josip S Herman, Dominic Grün, et al. Fateid infers cell fate bias in multipotent progenitors from single-cell rna-seq data. *Nature methods*, 15(5):379, 2018.
9. Dominic Grün et al. Revealing dynamics of gene expression variability in cell state space. *Nature methods*, 17:45–49, 2018.
10. Bo Wang, Junjie Zhu, Emma Pierson, Daniele Ramazzotti, and Serafim Batzoglou. Visualization and analysis of single-cell RNA-seq data by kernel-based similarity learning. *Nature Methods*, 14:414–416, 04 2017. doi: 10.1038/nmeth.4207.
11. Romain Lopez, Jeffrey Regier, Michael B Cole, Michael I Jordan, and Nir Yosef. Deep generative modeling for single-cell transcriptomics. *Nature methods*, 15(12):1053–1058, 2018.
12. Luyi Tian, Xueyi Dong, Saskia Freytag, Kim-Anh Lê Cao, Shian Su, Abolfazl JalalAbadi, Daniela Amann-Zalcenstein, Tom S Weber, Azadeh Seidi, Jafar S Jabbari, et al. Benchmarking single cell rna-sequencing analysis pipelines using mixture control experiments. *Nature methods*, 16(6):479–487, 2019.
